# DICAROS: Diffeomorphic Ancestral Shape Reconstruction on Phylogenies

**DOI:** 10.64898/2026.08.21.746152

**Authors:** Michael Lind Severinsen, Jacky Kaiyuan Li, Wonseop Lim, Levi Yoder Raskin, Gefan Yang, Stefan Sommer, Christy Anna Hipsley, Rasmus Nielsen

**Affiliations:** GeoGenetics Centre, Globe Institute, University of Copenhagen, Copenhagen 1350, Denmark; Biostatistics Division, University of California, Berkeley, CA 94720, USA; Department of Integrative Biology, University of California, Berkeley, CA 94720, USA; Department of Computer Science, University of Copenhagen, Copenhagen 1350, Denmark; Natural History Museum of Denmark, University of Copenhagen, Copenhagen 1353, Denmark; Department of Statistics, University of California, Berkeley, CA 94720, USA

## Abstract

**Summary:** Reconstructing ancestral morphologies on a phylogenetic tree is a central task in evolutionary morphometrics. Established reconstruction methods, including multivariate Brownian-motion approaches, rely on linear assumptions and do not directly model the correlations between landmarks within a shape, which can oversimplify the reconstructed morphology. The DICAROS method—Diffeomorphic Independent Contrasts for Ancestral Reconstruction of Shapes (Severinsen et al., 2026)— instead fuses sibling shapes along branches with large-deformation diffeomorphic (LDDMM) landmark dynamics that model these correlations, so that ancestors remain on the shape manifold. DICAROS was shown to outperform ordinary least-squares, Brownian-motion, and penalized-likelihood reconstruction, particularly on non-symmetric trees. `dicaros` repackages that pipeline as a documented, pip-installable tool that runs on arbitrary landmark datasets from a single command. It handles 2D and 3D landmarks, Newick and NEXUS trees, a choice of Euclidean or Fréchet species means, optional anchor-based alignment, and tips backed by a single specimen, and it returns the reconstructed shapes for all nodes together with the tree relabelled at its internal nodes. We demonstrate `dicaros` on two new datasets—a 2D leaf dataset (217 species) and a 3D guenon skull dataset (22 species).

**Availability and Implementation:** `dicaros` is a pip-installable Python package, freely available without registration under the MIT license at https://github.com/MichaelSev/DICAROS, with the two example datasets, documentation and reproduction scripts.

**Supplementary information:** Supplementary data (CPU and GPU runtime benchmarks) are available at *Bioinformatics* online.

## 1 Introduction

Geometric morphometrics summarizes biological form as configurations of homologous landmarks, and a recurring question is what the form of an ancestor looked like, given the forms of its descendants and a phylogeny (Bookstein, 1991; Adams and Collyer, 2019). Ancestral shapes are routinely reconstructed by fitting an evolutionary model to Procrustes-aligned landmark coordinates—by ordinary least squares, by maximum likelihood under Brownian motion, or by penalized likelihood (Slater et al., 2012; Adams and Otárola-Castillo, 2013; Revell, 2012)—and multivariate formulations explicitly account for the covariance among coordinates (Clavel et al., 2015). These approaches rely on linear assumptions in the Procrustes-aligned (Kendall) shape space and do not directly model the correlations between landmarks within a shape (Severinsen et al., 2026); even when among-coordinate covariance is included, the curved geometry of shape space is only linearly approximated, so reconstructed ancestral morphologies can be oversimplified.

A geometric alternative is to model the transition between two shapes as a smooth deformation (a diffeomorphism) and to reconstruct ancestors as shapes that lie on geodesics between their descendants, using the large-deformation diffeomorphic metric mapping (LDDMM) framework for landmarks (Joshi and Miller, 2000; Younes, 2010). Severinsen et al. (2026) implemented this as the DICAROS (Diffeomorphic Independent Contrasts for Ancestral Reconstruction of Shapes) method, which combines LDDMM landmark dynamics with Felsenstein’s phylogenetic independent contrasts (Felsenstein, 1985). When validated on swallowtail butterfly wings, DICAROS outperformed ordinary least-squares, Brownian-motion, and penalized-likelihood reconstruction in accuracy—particularly on non-symmetric trees, with comparable accuracy on symmetric ones. `dicaros` repackages the same pipeline as a documented, `pip`-installable tool that applies to arbitrary 2D and 3D landmark datasets from a single command, lowering the barrier to method application.

## 2 Methods

### Per-species means

Each individual specimen row is assigned to a species (a tip). Specimens within a species are Procrustes-superimposed (Gower, 1975; Rohlf and Slice, 1990) and reduced to a single mean shape, either the ordinary Euclidean mean or the Fréchet (Karcher) mean on the LDDMM landmark manifold; the species means are then aligned into one common frame. Alignment uses all landmarks by default, or only a user-supplied subset of anchor landmarks (`idxs`) when a few homologous points should define the frame. When a species is represented by a single specimen, that specimen is used directly as the tip mean, and a warning naming the tip is emitted, so the reconstruction proceeds rather than failing.

### DICAROS reconstruction

The reconstruction follows Severinsen et al. (2026): a single leaves-to-root message pass over a rooted, bifurcating tree (built on `hyperiax`; Yang et al., 2026). At each internal node, the two child shapes are fused on the landmark manifold (using `jaxgeometry`; Sommer and Kühnel, 2021): the shorter child branch defines a base point, the Riemannian logarithm toward the other child gives a momentum (a diffeomorphic independent contrast), and the Hamiltonian geodesic is integrated to the point along the branch fixed by the relative branch lengths. The fused estimate becomes the node’s shape, and the pass continues to the root, so every ancestor is itself a valid shape on the landmark manifold rather than a linear interpolation of coordinates. In `dicaros`, the Gaussian kernel width *σ* that sets the spatial correlation between landmarks is taken from the mean nearest-neighbor landmark spacing, so it matches the data scale.

### Inputs and outputs

Landmark tables are plain CSV; coordinate columns are selected by regex, start index, or explicit list, accommodating arbitrary leading metadata columns. Trees may be Newick or NEXUS, including 10kTrees-style `translate` tables (Arnold et al., 2010), and are pruned to the species present in the data, with polytomies arbitrarily resolved with minuscule branch lengths. The output is (i) a CSV of reconstructed landmark configurations for *all* nodes (tips and ancestors), and (ii) the input tree with internal nodes labeled, so each CSV row maps back onto the phylogeny. Implementation is in Python on top of JAX (Bradbury et al., 2018); the diffeomorphic step runs on CPU by default.

## 3 Usage and examples

From the command line:

~~~
dicaros --landmarks leaves.csv --tree tree.nwk \
        --dim 2 --species-col species \
        --landmark-regex ‘^[xy][0-9]+$’ \
        --output-dir out/leaf
~~~

or through the `dicaros.reconstruct(…)`function equivalently. The user chooses the mean estimator (`-mean euclidean|frechet`) and whether to anchor alignment on specific landmarks (`-idxs`).

Whereas DICAROS was originally validated on swallow-tail butterfly wings (Severinsen et al., 2026), we applied `dicaros` to two further, contrasting datasets (Table 1): a 2D dataset of 102-landmark grass (Poaceae) leaf outlines for 217 species (Gallaher et al., 2019a,b), and a 3D dataset of 155-landmark guenon skulls for 22 species (Cardini and Elton, 2017) on the 10kTrees primate phylogeny (Arnold et al., 2010). The leaf dataset includes 17 single-specimen tips, and the guenon dataset includes one, all handled automatically. For each dataset `dicaros` reconstructs a shape at every ancestral node; Figure 1 shows the resulting phylomorphospaces (a principal-component ordination of all node shapes with the tree edges drawn in shape space) and the reconstructed root overlaid on the tip means. ^1^

**Table 1.**
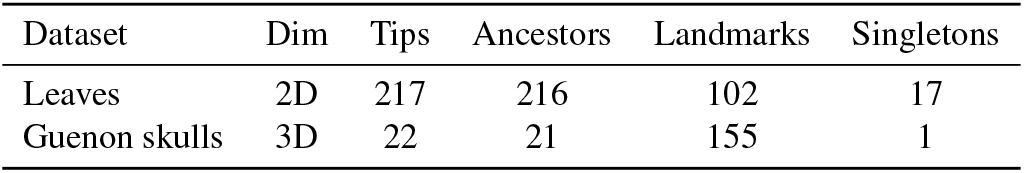
Reconstruction on the two bundled datasets. Ancestors = internal nodes reconstructed; singletons = tips backed by one specimen.

**Figure 1.**
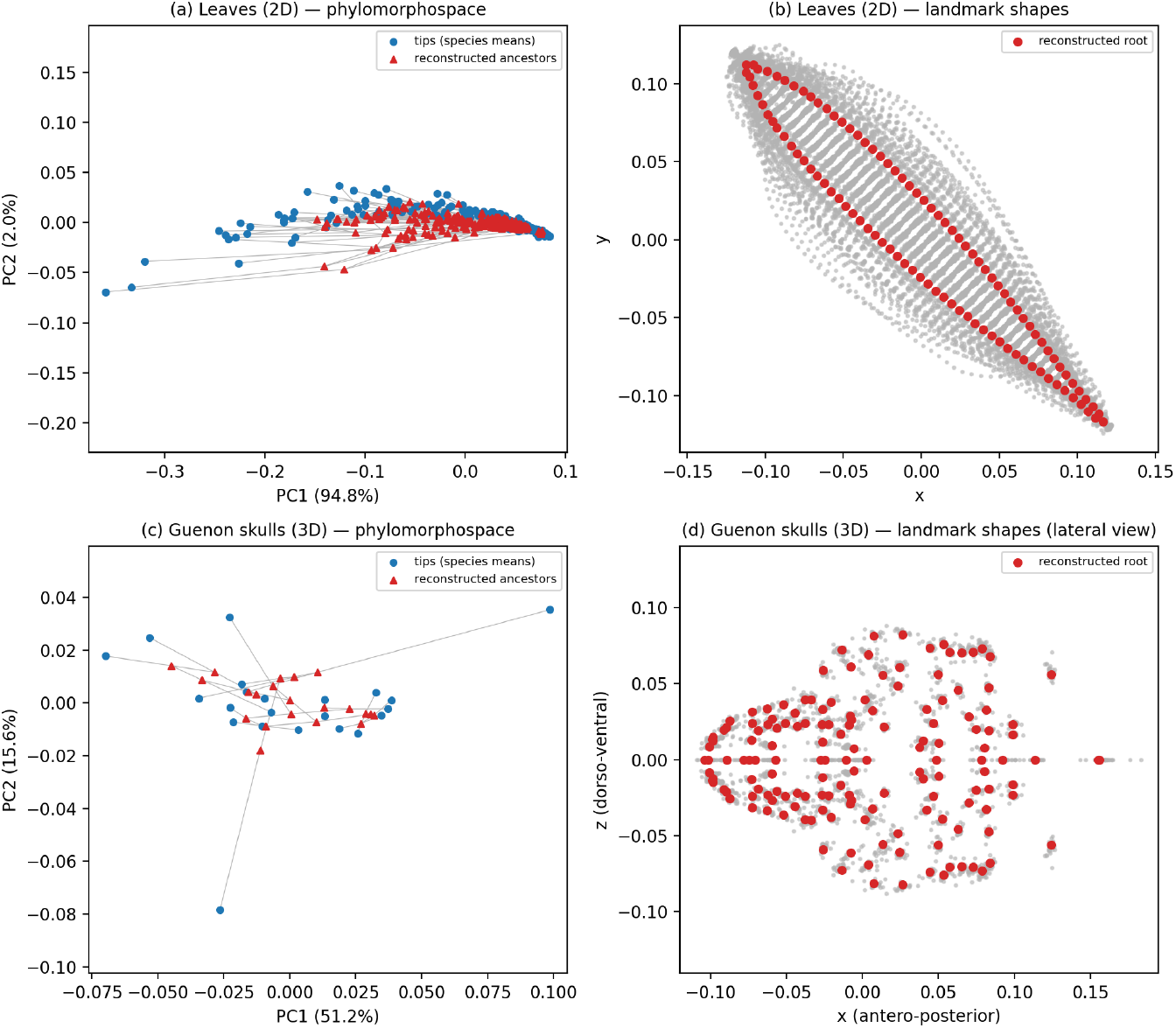
Reconstruction on the two example datasets. Top, leaf dataset (2D, 217 species); bottom, guenon skull dataset (3D, 22 species). (a,c) Phylomorphospace: a principal-component ordination of all node shapes (blue, tip means; red, reconstructed ancestors) with the tree edges drawn in shape space. (b,d) Reconstructed root configuration (red) overlaid on the tip mean shapes (grey); the 3D guenon skulls are shown in lateral view (the raw *x, z* plane of the Procrustes-aligned coordinates: antero-posterior *×* dorso-ventral).

During the ongoing maintenance of the package we plan to update `dicaros` continuously and add new features as extensions of the current methodology. One planned direction is to adopt landmark shape spaces whose induced metric descends from a right-invariant Sobolev metric on the diffeomorphism group and is itself invariant to rigid motions and scale (Joshi et al., 2026), so that Procrustes pre-alignment and diffeomorphic deformation are handled by a single geometry rather than as separate steps, while retaining the regularity that keeps landmarks from colliding.

## Acknowledgements

We thank the developers and maintainers of `hyperiax` and `jaxgeometry`.

## Funding

This work was supported by a research grant (VIL40582) from VILLUM FONDEN.

## Conflict of Interest

None declared.

## Supplementary Material

**Table S1.** Reconstruction wall-clock time for dicaros on the two example datasets, on CPU and GPU. CPU runs were executed on an Intel Xeon Gold 5420+, pinned to 32 logical cores; the GPU runs used a single NVIDIA RTX 6000 Ada Generation (49 GB). Times are wall-clock and include JAX just-in-time compilation. CPU and GPU reconstructions produce the same tree topology and internal-node labels, and agree on node coordinates to floating-point reduction tolerance (guenon: maximum coordinate difference 2.9 *×* 10^−4^; leaves: maximum 1.3 *×* 10^−2^, mean 2.0 *×* 10^−5^).

| Dataset | Landmarks (dim) | Tips | CPU time (32 cores) | GPU time (RTX 6000 Ada) |
| --- | --- | --- | --- | --- |
| Leaves | 102 (2D) | 217 | 1.70 h (102 min) | 10.2 min (609 s) |
| Guenon skulls | 155 (3D) | 22 | 0.22 h (13 min) | 2.1 min (124 s) |

Runs used `dicaros` v0.1.0 with `hyperiax` 3.0 and JAX. The DICAROS ancestral reconstruction (Hamiltonian geodesic integration with the flow-differential lift) is the dominant cost; both the per-species mean-shape estimation and the tree handling are negligible by comparison. The CPU figures are a conservative reference point (32 of 112 available cores); GPU execution is roughly 6–8 × faster on these datasets.

## Footnotes

1 For illustration, an animation of the reconstructed guenon skull (lateral view) morphing along a root-to-tip lineage is available in the project repository (https://github.com/MichaelSev/DICAROS). It is a standalone visualization, generated with the notebook at https://github.com/MichaelSev/DICAROS/blob/DICAROS-Publication-Systematic-Biology/Image_Trajectory.ipynb, and is not part of the DICAROS package.

## Notes

### Competing Interest Statement

The authors have declared no competing interest.

